# A High-fat, High-salt Diet Model of MDAKD Impairs Bioenergetic Efficiency for ATP Synthesis

**DOI:** 10.64898/2026.02.20.707069

**Authors:** Stephen T Decker, Za’rya T Smith, Precious C Opurum, Venisia L Paula, Kennedy N Moses, Deborah Stuart, Anu S Kurian, Subhasmita Rout, Nirupama Ramkumar, Katsuhiko Funai

## Abstract

Metabolic dysfunction-associated kidney disease (MDAKD) is closely linked to dietary excess, but models that capture early kidney injury without obesity are limited. We fed male C57BL/6J (6J) and C57BL/6N (6N) mice a high-fat, high-sodium (HF/HNa) or control diet for 16 weeks. HF/HNa feeding did not alter body weight, adiposity, or total food intake; however, it increased dietary energy and sodium exposure, kidney mass, water intake, and urine volume. GFR declined modestly in 6J mice, whereas 6N mice maintained or slightly increased GFR. Both substrains showed increased urinary albumin, creatinine, KIM-1, and NGAL, while cystatin C rose predominantly in 6N mice, indicating strain-dependent tubular injury. Whole-kidney trichrome staining revealed increased fibrotic area with HF/HNa, particularly in 6N mice, without significant changes in glomerular morphology. In isolated renal mitochondria, oxygen consumption was preserved, but ATP production and ATP:O ratios were reduced, with unchanged citrate synthase activity and OXPHOS protein abundance, consistent with early mitochondrial bioenergetic uncoupling. Exploratory urinary proteomics in 6J mice identified HF/HNa-associated changes in proteins linked to tubular stress and extracellular matrix remodeling. These findings define an early MDAKD-like renal phenotype with strain-specific functional responses, tubular injury, fibrosis, and impaired mitochondrial ATP efficiency.

**Translational Statement:** Metabolic Dysfunction-Associated Kidney Disease (MDAKD) is a leading driver of chronic kidney disease (CKD) in the world. In addition to obesity and related comorbidities, renal mitochondrial dysfunction is thought to be a key contributor to the development of CKD in patients with MDAKD; however, few models recapitulate the progression of MDAKD. We couple well-established mouse models of obesity, namely the C57Bl/6J and C57Bl/6N mouse lines, with a high-fat, high-salt diet to induce renal mitochondrial dysfunction, leading to early stages of MDAKD as indicated by widespread fibrosis and mild reduction in glomerular filtration rate, though these effects were strain-dependent. We identify diet-induced mitochondrial dysfunction as a common feature in both mouse strains, suggesting impairments in mitochondrial respiration and oxidative ATP production are indeed a contributing factor to the development of MDAKD. This study highlights the role of energetic impairments in the pathogenesis of MDAKD and may guide future therapies for CKD.

## Introduction

Chronic kidney disease (CKD) is a major global health burden, affecting nearly 10% of the population worldwide and accounting for more than 800 million individuals overall.^1,2^ CKD is strongly associated with increased risk of cardiovascular disease and premature mortality, with the majority of patients dying from cardiovascular complications rather than progressing to kidney failure.^1,3,4^ One of the dominant drivers of CKD in contemporary populations is the metabolic dysfunction-associated kidney disease (MDAKD), a pathophysiological state defined by the convergence of obesity, metabolic dysregulation, and renal dysfunction.^3–5^ Patients with MDAKD exhibit accelerated loss of glomerular filtration rate (GFR), higher albuminuria, lipid deposition in the kidney, and a greater risk of progression to end-stage kidney disease compared to individuals with isolated kidney injury.^5,6^

Despite its prevalence and clinical importance, preclinical models that recapitulate the multifactorial nature of MDAKD-associated CKD remain lacking.^7^ Traditional murine models of CKD rely on surgical interventions such as unilateral ureteral obstruction or 5/6 nephrectomy, which produce acute and severe injury but fail to mimic the gradual, systemic progression of human disease.^8^ Genetic models replicate rare monogenic disorders but lack relevance to the common multifactorial pathogenesis of MDAKD.^9^ Dietary manipulations such as high-fat feeding^10–17^, which induce obesity and insulin resistance, and high-salt diets, which promote hypertension and tubular stress^10,12,18,19^, have been widely used. However, in isolation, these approaches rarely lead to measurable declines in GFR or sustained CKD progression. Thus, no standardized diet-focused murine paradigm faithfully captures the integration of systemic metabolic load, tubular injury, and mitochondrial dysfunction that typifies human MDAKD-associated CKD.^3,8,20–22^

There is increasing recognition that impaired renal mitochondrial respiration and ATP synthesis are a central mediator of progressive CKD, particularly in the setting of increased tubular sodium reabsorption and metabolic overload.^16,21,23–29^ The kidney is among the most energy-demanding organs, with sodium reabsorption accounting for the majority of ATP utilization.^22^ Maladaptive increases in transport work, particularly in the proximal tubule and thick ascending limb, can impose a metabolic burden that exceeds mitochondrial capacity, leading to oxidative injury, fibrosis, and nephron loss.^13,16,21,22^ However, few murine studies have systematically linked dietary stressors with renal functional decline, structural injury, transporter adaptations, and mitochondrial bioenergetics in vivo. Furthermore, although reports have suggested that standard mouse models of obesity, such as the C57Bl/6J and C57Bl/6N strains, are less prone to kidney disease,^30^ the combination of a high-fat, high-salt diet in these strains has not yet been utilized in a renal-specific context.

In this study, we tested whether long-term exposure (16 weeks) to a combined high-fat, high-salt (HF/HNa) diet could reproduce key renal features of MDAKD. We evaluated systemic and renal physiology, histopathology, injury marker expression, and mitochondrial respiration using a substrate–uncoupler–inhibitor titration (SUIT) protocol.^31^ We aimed to determine if HF/HNa feeding recapitulates traits of human CKD, and whether this approach could serve as a relevant model for future mechanistic and therapeutic research.

## Methods

### Animal Care & Husbandry

8-10-week-old male C57BL/6J mice (Strain No. 000664; n = 23) were purchased from Jackson Laboratory, and C57BL/6NCrl (Strain code: 027; n = 28) mice were purchased from Charles River. Figure 1A shows a diagram of the experimental approach. All mice were maintained in our colony and randomly assigned to a standard chow diet or a 42% fat, ∼8% salt diet (HF/HNa; Envigo Custom TD.230162; Figure 1B) for 16 weeks. Mice were housed in standard conditions, and all animals had access to *ad libitum* food and water throughout the study duration. Body weights were measured weekly. End-of-study body composition was determined using a Bruker Minispec NMR (Bruker, Billerica, MA). All animals were handled according to approved University of Utah Animal Use and Care Committee (Protocol No. 1633).

**Figure 1.**
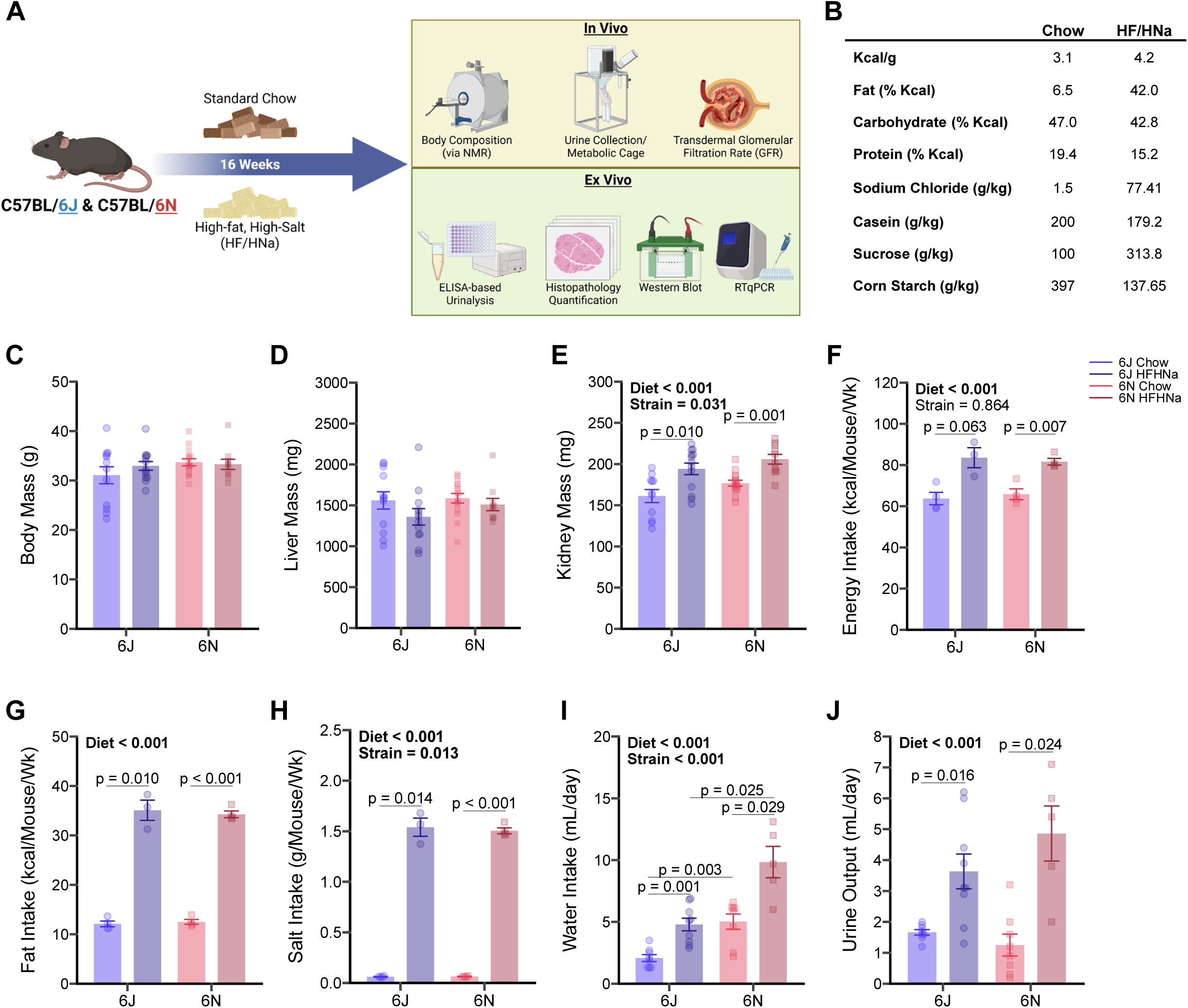
Experimental setup (A) and composition of the diets (B) of the present study. Body weight (C) and liver mass (D) were not different between groups at the time of termination; however, kidney mass (E) was significantly greater in high-fat, high-salt (HFHNa) fed mice compared to their respective controls. Free-living, per-cage energy intake (F) was greater in the HFHNa-fed mice than in the chow-fed mice, as was fat (G), salt (H), and water (I) intake. Individual urine output (J) collected during a 24-hour period in metabolic cages was also increased in the HFHNa-fed mice.

### Glomerular Filtration Rate (GFR)

Transcutaneous GFR was measured using the clearance of fluorescein isothiocyanate-labeled sinistrin (FITC-sinistrin) with a commercially available device (NIC-Kidney, MediBeacon GmbH, Mannheim, Germany), as previously described,^32^ according to the manufacturer. Mice were placed under non-lethal isoflurane-induced anesthesia. A patch of dorsal skin was shaved, and the monitor was attached. Background fluorescence was measured for 10 min, after which FITC-sinistrin (7.5 mg/100 g body wt) was administered retro-orbitally. FITC-sinistrin fluorescence was monitored for the next 90 minutes. The kinetics of fluorescence disappearance were used to determine GFR using the manufacturer’s software, according to their standard guidelines. We measured the GFR of all mice between 10:00 AM and 02:00 PM.

### Urine Collection & Urinalysis

Mice were placed in an MMC100 Metabolic Cage, and urine was collected every 24 h for 48 h at the end of the study. Body weight, food, and water intake were all observed throughout the experiment. Mice were housed with their respective chow or HF/HNa diet converted into a gel with free access to water. Urine samples were centrifuged at 21,000 g for 15 min, then aliquoted and stored at −80°C. Urine albumin and creatinine were assessed using an absorbance-based Albumin Creatinine Ratio Assay Kit (ab241018; Abcam Ltd., Cambridge, UK). Urine Cystatin C (CysC; kit KE10066), lipocalin 2 (Lcn2, or NGAL; kit KE10045), and kidney injury marker-1 (KIM-1, or HAVCR1; kit KE10064) were assessed using commercially available kits from Proteintech (Rosemont, IL, USA). All assays were performed according to the manufacturer’s recommendations.

### Euthanasia, Organ Harvesting, and Blood Plasma Collection and Analysis

After 16 weeks of the dietary intervention, mice were fasted for 2 hours (beginning at 7:00-8:00 AM to minimize feeding artifacts and variability in metabolic activity caused by food intake and circadian rhythm) before blood was collected, and animals were euthanized using a lethal mix of ketamine and xylazine. Blood was collected in EDTA-coated blood collection tubes and centrifuged at 1,000 x g for 10 minutes. The separated plasma was then aliquoted and stored at −80°C for later analysis. Blood urea nitrogen (BUN) was analyzed using the QuantiChrom^TM^ Urea Assay Kit (DIUR-100; BioAssay Systems, Hayward, CA, USA) according to the manufacturer’s protocol.

Following euthanasia, the kidneys were then harvested and weighed. At random, one kidney was finely cut into thirds, snap-frozen in liquid nitrogen, and stored for later western blotting and qPCR. The other kidney was finely sliced for histological staining and mitochondrial function assays.

### Histology

Kidney samples were fixed in 10% formaldehyde for 24–48 h and transferred to the ARUP Research Histology Core Laboratory at the University of Utah. Kidney slices were fixed in paraffin, sectioned at 4 μm, and stained with H&E, PAS, and Masson’s trichrome to detect morphological changes and fibrosis. Trichrome-stained kidney sections were imaged using Zeiss AxioScan.Z1 (ZEISS, Oberkochen, Germany) slide scanner at the University of Utah Imaging core.

### Histology Analysis

Trichrome-stained kidney sections were analyzed in Fiji software^33^ (version 2.7.0) using a combination of custom macros, the Labkit plugin^34^ (version 0.4.0), and a custom-made Python script to segment images using the MONAI framework^35–37^ in a Google Colab environment, and are described in detail in the supplementary methods. Briefly, glomerular and tubular structures were identified to generate mask image files for all slides. The generated masks were used to crop individual glomeruli and tubular structures for analysis. Features were analyzed following postprocessing feature cutoffs and the colour deconvolution2^38–40^ (version 2.1) plugin incorporated into a custom macro in Fiji.

### Kidney Mitochondrial Isolation

Mitochondria isolation was performed as previously described.^41^ Briefly, kidney sections were minced in ice-cold mitochondrial isolation medium (MIM) buffer (300 mM sucrose, 10 mM HEPES, 1 mM EGTA; pH 7.4) with 1 mg/mL bovine serum albumin (BSA)and carefully homogenized with a Teflon-glass pestle. The homogenate was centrifuged at 800 g for 10 min at 4°C. The resulting supernatant was transferred to another tube and centrifuged at 1,300 g for 10 min. The resulting supernatant was transferred to another tube and centrifuged at 10,000 g for 10 min at 4°C. The pellet was then resuspended in 100-300 μL of MIM buffer. Protein concentrations were assessed with Pierce BCA Protein Assay Kits (No. 23225, Thermo Fisher Scientific, Waltham, MA, USA).

### High-Resolution Respirometry and Fluorometry

Mitochondrial O_2_ consumption (*J*O_2_) was evaluated using the Oroboros Oxygraph O2K (Oroboros Instruments, Innsbruck, Austria), as previously described.^41^ Isolated mitochondria were introduced to the oxygraphy chambers containing 2 mL of Buffer Z [105 mM MES potassium salt, 30 mM potassium chloride (KCl), 10 mM monopotassium phosphate (KH2PO4), 5 mM magnesium chloride (MgCl2), and 0.5 mg/mL BSA]. Respiration was assessed by adding the following substrates: 0.5 mM malate (M), 5 mM pyruvate (P), 5 mM glutamate (G), 2 mM ADP, 10 mM succinate (S), and 1.5 μM carbonyl cyanide-p-trifluoromethoxyphenylhydrazone (FCCP).

ATP production (*J*ATP) was fluorometrically quantified with a Horiba Fluoromax 4 (Horiba Scientific, Piscataway, NJ, USA) by enzymatically linking it to NADPH synthesis, as previously described.^41^ ATP generation was assessed with 0.5 mM malate, 5 mM pyruvate, 5 mM glutamate, 10 mM succinate, and successive 2, 20, and 200 μM ADP additions.

### Western Blot Analyses

Whole kidney tissue samples were homogenized in cold RIPA lysis buffer (No. 89901, Thermo Fisher Scientific) containing protease inhibitor (No. 78446, Thermo Fisher Scientific), incubated at 4°C for 10 min, and centrifuged at 12,000 g for 15 min at 4°C, and the supernatant was collected into a fresh tube. The BCA Protein Assay Kit (No. 23225, Thermo Fisher Scientific) was used to measure the protein concentration in the supernatant. Western blotting was carried out as previously described^41^ with slight modifications. In summary, for electrophoresis, appropriate volumes of protein were combined with 4× Laemmli sample buffer supplemented with 10% β-mercaptoethanol (#No. 1610710, Bio-Rad, Hercules, CA) and loaded onto a Bio-Rad 4%–20% gradient SDS-polyacrylamide gel. After that, the proteins were transferred from the gel onto membranes made of polyvinylidene fluoride. The membranes were blocked for 1 h at room temperature using 5% low-fat skim milk in Tris-buffered saline containing 0.1% Tween 20 (TBS-T), and then, the kidney samples were incubated overnight with specific primary antibodies (Table 1) to probe for protein abundance. Following multiple TBS-T washes, the membranes were incubated for 1 h at room temperature with appropriate secondary antibodies diluted 1:5,000 in 5% BSA. After multiple washes in TBS-T and one final rinse in TBS. Membranes were exposed with Western Lightning Plus-ECL (PerkinElmer) and imaged using a ChemiDoc Imaging System (Bio-Rad, Hercules, CA). The images were quantified using Image Lab software (Bio-Rad).

### Real-Time Quantitative Polymerase Chain Reaction (qPCR)

For qPCR investigations, whole mouse kidney tissues were lysed in 1 mL of TRIzol (Thermo Fisher Scientific), and RNA was extracted using conventional methods. The iScript cDNA Synthesis Kit (Bio-Rad, Hercules, CA) was used to reverse-transcribe total RNA. To assess the expression of kidney damage biomarkers (Table 1). Prevalidated primer sequences were taken from either the primer bank or previously published papers. All mRNA levels were standardized to RPL32 with SYBR Green reagents (Thermo Fisher Scientific), and 384-well plates were loaded and sent to the University of Utah Genomics Core laboratories for processing.

### Mass Spectrometry-based Proteomics

Mass spectrometry–based proteomics was performed as described in detail in the supplementary methods. Briefly, urinary proteins (∼200 μL) were acetone-precipitated, processed using S-TRAP columns with reduction, alkylation, and trypsin digestion, and peptides were eluted, desalted, and resuspended in formic acid; a pooled QC sample and a fractionated pooled library were generated. Peptides (200 ng) were analyzed by nano–LC-MS/MS on a nanoElute 2 coupled to a Bruker timsTOF Pro2 using a 45-min reversed-phase gradient, with PASEF-DDA for library generation and dia-PASEF for sample analysis. A custom spectral library was built in FragPipe v22.0 against the UniProt mouse reference proteome, and protein quantification was performed in DIA-NN v2.0 with standard variable modifications, one missed cleavage allowed, and a 1% false discovery rate at the peptide and protein levels.

### Statistical Analysis

All statistical analyses were performed in R (version 4.5.1). Data are reported as mean ± SEM unless otherwise noted. Group comparisons were analyzed using n-way ANOVA with diet (chow vs. HF/HNa) and strain (C57Bl/6J vs. C57Bl/6N) as fixed factors. Normality and variance assumptions were evaluated, and nonparametric aligned rank transform^42,43^ (ART) procedures were applied when appropriate. Post hoc multiple comparisons were performed using the Holm–Šidák method. For histological segmentation and glomerular morphometry, linear mixed-effects models (LMM) were employed with individual mice as a random factor to account for nested sampling. For urine proteomics analysis, data were processed as described above, and significance was corrected for the False Discovery Rate (FDR). Analyses were conducted using the R packages *tidyverse, ggprism, ggbeeswarm, rstatix, ARTool, ggpubr, multcomp, reshape2, ggcompare, mutoss, emmeans,* and *svglite*. Statistical significance was defined as p < 0.05.

## Results

### HF/HNa feeding alters fluid balance and intake without changing body weight or adiposity

Over the 16-week intervention (Figures 1A & 1B), both C57Bl/6J (chow n = 13, HF/HNa n = 10) and C57Bl/6N mice (chow n = 15, HF/HNa n = 13) gained weight progressively, but endpoint body weight and liver mass did not differ by diet or strain (Figures 1C & 1D). However, kidney mass was increased in the HF/HNa diet for both strains (Figure 1E), indicating that the combined high-fat, high-salt diet did not induce obesity or gross changes in adiposity compared with chow controls (likely due to no differences in food overall intake, shown in Supplementary Figure S1), but did induce renal hypertrophy. As expected, mice on the HF/HNa diet had a greater energy, fat, and salt intake compared to the chow groups (all p < 0.001; Figures 1F-H), consistent with a sustained shift in osmotic balance. In parallel, diet increased water intake (p < 0.01; Figure 1I) and urine output (p < 0.001; Figure 1J). Body composition was not influenced by diet in either strain (Supplementary Figure S1).

### Renal Function

GFR, assessed by transcutaneous FITC–sinistrin clearance, differed by diet and substrain (Figures 2A–B). In C57Bl/6J mice, HF/HNa feeding significantly reduced GFR compared with chow controls, whereas in C57Bl/6N mice, HF/HNa-fed animals showed a small, non-significant increase in GFR relative to chow.

**Figure 2.**
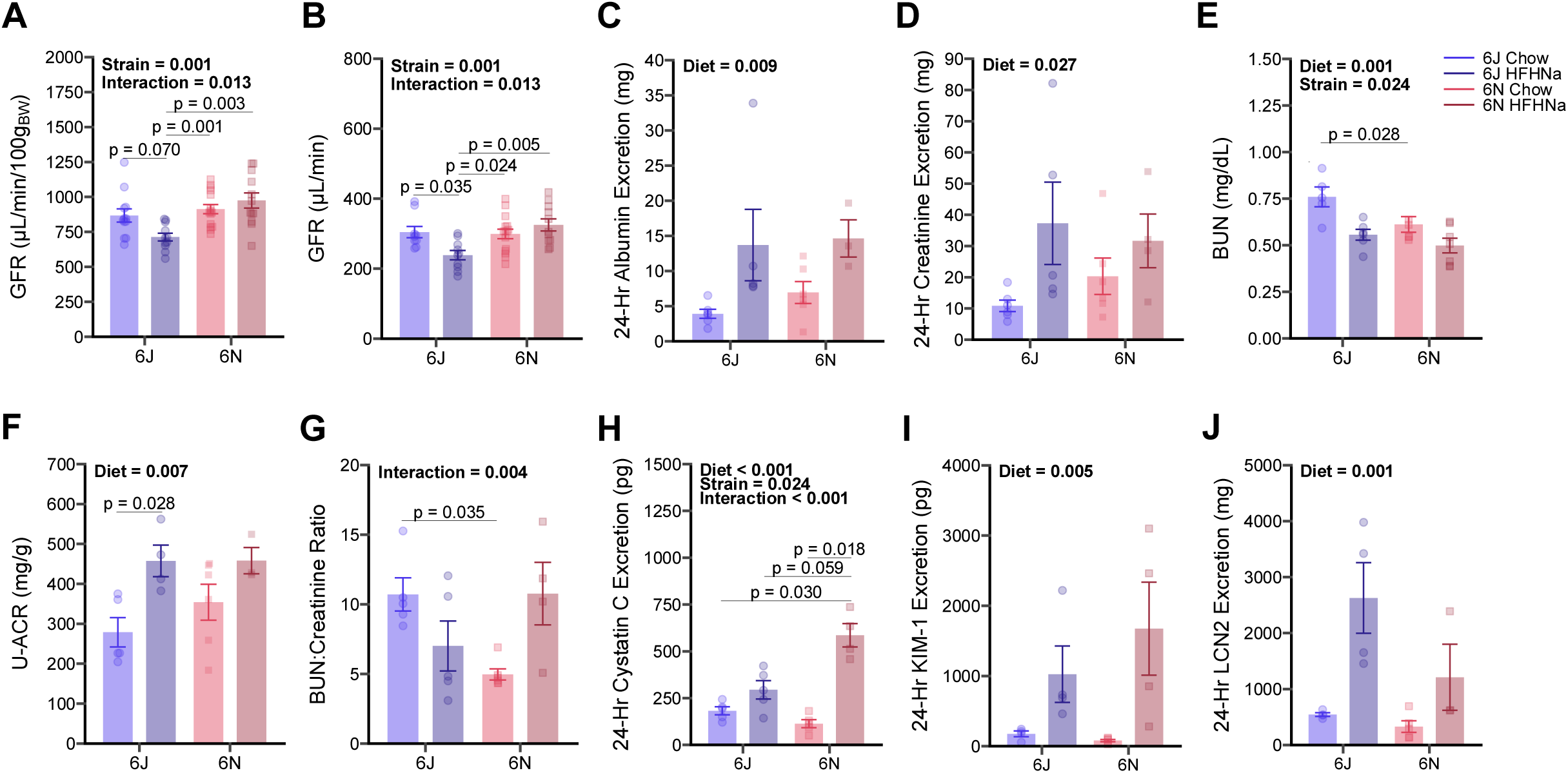
Glomerular filtration rate (GFR; A & B) was mildly (∼20%) decreased in the 6J high-fat, high-salt-fed (HFHNa) compared with chow-fed mice, but no differences between diet groups were observed in 6N mice. However, 24-hour urine albumin (C), creatinine (D), and albumin were significantly increased in the HFHNa-fed mice, while plasma blood urea nitrogen (BUN; F) was significantly decreased. Albumin-to-creatinine ratio (ACR; F) was likewise significantly increased in the 6J strain, but there were no significant differences in the BUN-to-creatinine ratio (G). In agreement with other markers of impaired kidney function, urine cystatin C (H), kidney injury marker 1 (KIM-1; I), and lipocalin 2 (LCN2; J) were increased by HFHNa feeding.

HF/HNa feeding increased urinary albumin and creatinine excretion and decreased plasma BUN in both substrains (Figures 2C–E). Urinary albumin–creatinine ratio (ACR) showed a main effect of diet and was significantly higher in HF/HNa-fed C57Bl/6J mice, with more variable responses in C57Bl/6N mice (Figure 2F). BUN:creatinine ratios did not differ significantly between diets in either substrain (Figure 2G). Urinary cystatin C excretion was significantly higher in HF/HNa-fed C57Bl/6N mice but not in C57Bl/6J mice (Figure 2H). Urinary KIM-1 and LCN2 concentrations were also higher in HF/HNa-fed animals, with a significant main effect of diet (p = 0.005 and p = 0.001, respectively; Figures 2I–J).

### Histopathological Analysis

Structural analysis of glomeruli revealed significant remodeling with HF/HNa feeding. Representative Masson’s trichrome staining showed increased collagen deposition in HF/HNa kidneys compared with chow (Figure 3A; representative images of H&E and PAS stains are shown in Supplemental Figure 3). Whole-slide kidney fibrosis stain area was increased in the HF/HNa-fed animals (main effect p < 0.001; Figure 3B). However, 6N showed a much greater increase in fibrotic area relative to their chow counterparts (Chow vs HFHNa: p < 0.001) than the 6J mice (Chow vs HFHNa: p = ns).

**Figure 3.**
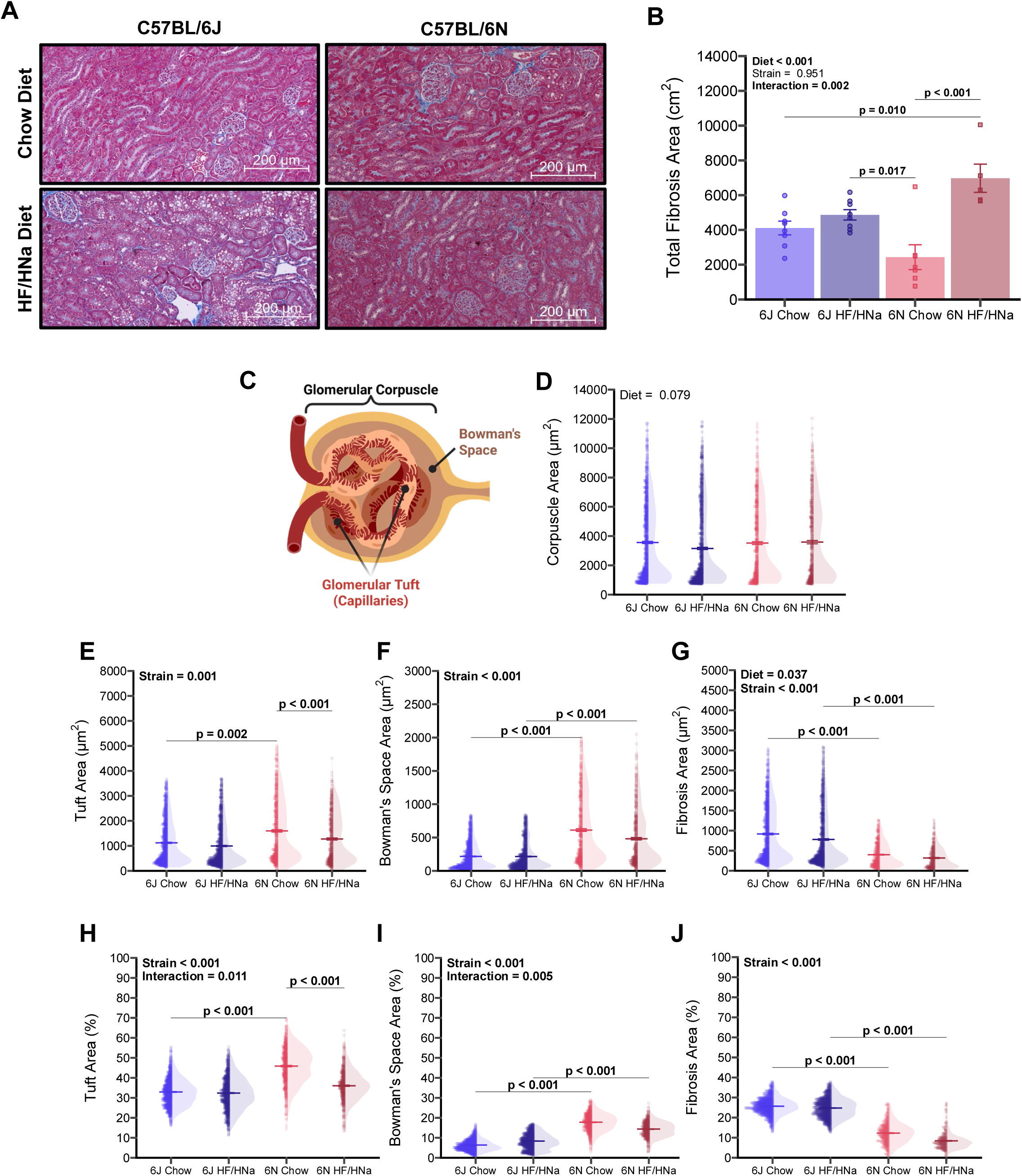
A representative image of Masson’s trichrome stain is shown in panel A. A high-fat, high-salt diet (HFHNa) significantly increased fibrosis area in the 6N, but not 6J, mice. Glomerular-specific analysis of the glomerular corpuscle, tuft, Bowman’s space, and fibrosis areas was then assessed (C). There was no significant difference in corpuscle area (D), tuft area (E & H), or Bowman’s space area (F & I). However, HFHNa diet decreased the absolute (G) but not the relative (J) fibrosis stain area.

A custom Python pipeline was used to segment (representative image shown in Supplementary Figure 3A) and generate binary masks of individual glomeruli, which were subsequently filtered by size and circularity to exclude artifacts. A schematic of the glomerular compartments, highlighting the features subjected to quantification, is shown in Figure 3C. Quantitative analysis following color deconvolution revealed multiple abnormalities in glomerular structure. A HF/HNa diet did not change total corpuscle area (Figure 3D), nor did it change tuft or Bowman’s Space areas (Figures 3E, 3F, 3H, and 3I), but it did significantly decrease glomerular fibrosis area (p < 0.05; Figure 3G and 3J). We did not observe any significant changes in the fibrotic areas of the tubules or in the basement membrane, nor did we see substantial changes in Col1α1 or Col3α1 mRNA (Supplementary Figure 3). Taken together with the functional data, these findings suggest that the HF/HNa-associated decline in GFR observed in 6J mice likely reflects widespread tubular impairments and/or altered hemodynamics rather than established glomerulosclerosis or tubulointerstitial fibrosis at this stage.

### Transporter and Inflammatory Protein Expression

Western blot analysis revealed evidence of strain-dependent responses of renal transporters and podocyte proteins to HF/HNa feeding (representative blots shown in Figure 4A). In 6N mice, AQP2 abundance was increased (p = 0.051; Figure 4B), whereas 6J mice showed no change compared with chow controls. NKCC2 abundance was unchanged in both strains (Figure 4C). However, SGLT2 abundance trended toward a decrease in 6N mice on HF/HNa (p = 0.062; Figure 4D), but not in 6J mice, suggesting adaptation to sodium reabsorptive stress is a feature unique to 6N mice. Further, the mechanism underlying these adaptations is not the result of changes in ER stress, as the ER stress markers BiP, CHOP, and PERK were unchanged (Figures 4E-4G). Podocin abundance was unchanged in both strains (Supplemental Figure 4B), suggesting minimal podocyte injury in these models. Likewise, Na/K^+^-ATPase and TNF-α abundance were also unaffected by diet (Supplemental Figure 4C and 4D).

**Figure 4.**
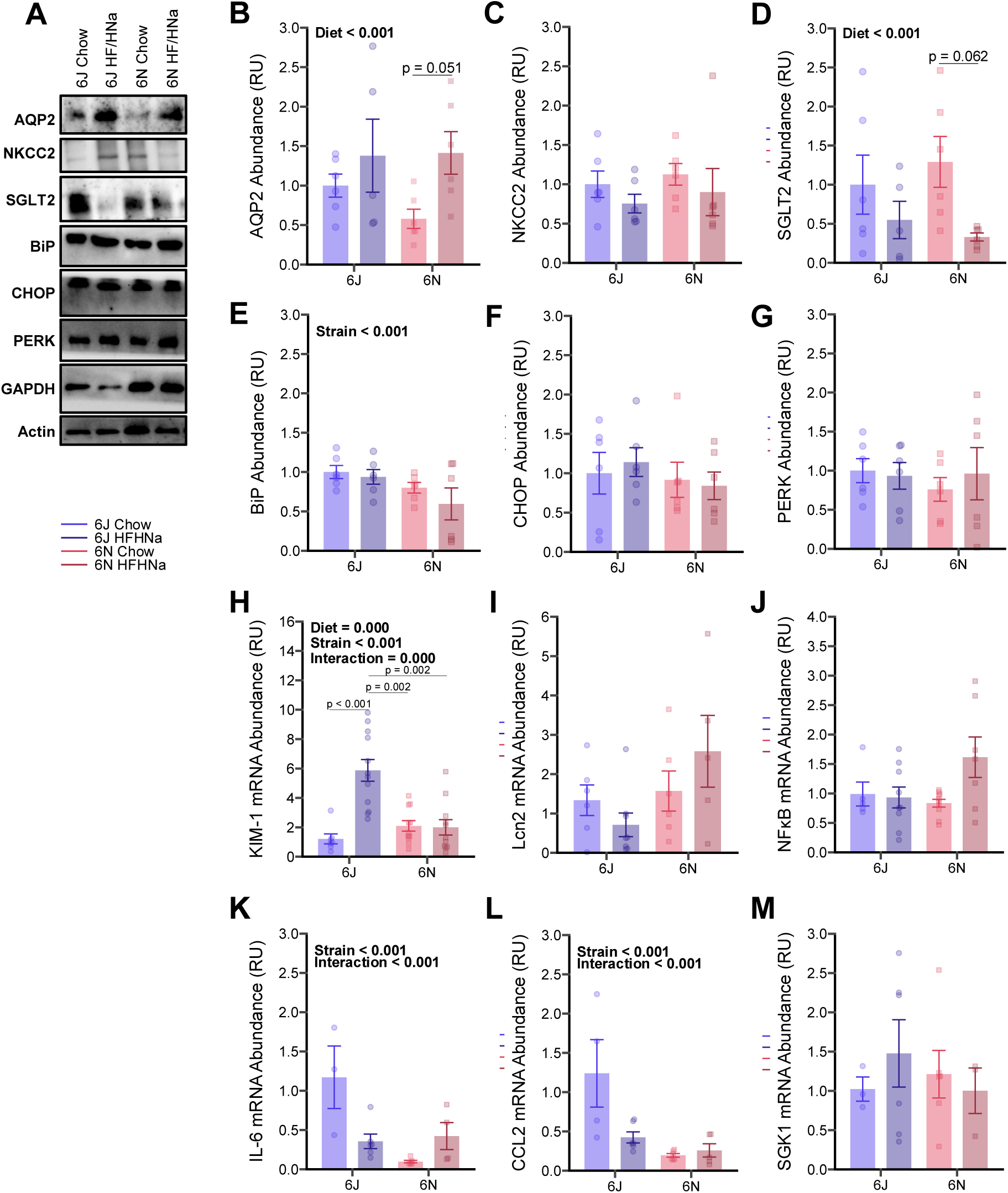
Representative images of western blots are shown in panel A. A high-fat, high-salt diet (HFHNa) resulted in a significantly increased protein abundance of aquaporin 2 (AQP2; B), but had no effect on sodium-potassium cotransporter 2 (NKCC2; C). On the other hand, HFHNa feeding resulted in a significant decrease in the protein abundance of the sodium-glucose cotransporter (SGLT2; D). No changes were observed in the protein abundance of the ER stress markers BIP (E), CHOP (F), or PERK (G). mRNA abundance of kidney injury marker (KIM-1; H) was increased in only the 6J HFHNa-fed mice. Diet did not significantly alter the mRNA abundance of lipocalin 2 (LCN2; I), NFκB (J), interleukin-6 (IL-6; K), C-C motif chemokine ligand 2 (CCL2; L), or serum and glucocorticoid-induced protein kinase 1 (SGK1; M).

At the mRNA level, 6J, but not 6N, mice fed a HF/HNa diet exhibited significantly increased mRNA abundance of KIM-1 (p < 0.001; Figure 4H). Surprisingly, diet did not induce any changes in Lcn2, NFκB, or any canonical markers of inflammation (Figures 4I-M). Together, these findings suggest that 6J mice experience a greater burden of osmotic imbalance under HF/HNa feeding, which results in tubular injury that is not dependent on ER stress or inflammatory pathways. Alternatively, the adaptive responses in 6N kidneys may underlie their preserved or slightly elevated GFR despite similar exposure to HF/HNa feeding.

### Mitochondrial Respiration and ATP Production

Kidney cells (especially proximal tubules) are highly energetic and rely heavily on mitochondrial oxidative phosphorylation to maintain the reabsorptive load placed on the kidneys, and thus, global osmotic balance. High-resolution respirometry of isolated kidney mitochondria showed diet-linked differences in mitochondrial function (Figure 5A–D). While the 6N mice exhibited constantly higher *J*O_2_ compared to 6J mice, HF/HNa feeding did not alter mitochondrial respiration under any of the substrate conditions. However, the HF/HNa diet did significantly impair *J*ATP and, thus, lowered the ATP:O ratio, indicating an uncoupling effect and possible bioenergetic impairment of HF/HNa diet on kidney mitochondria. Furthermore, it is unlikely that these changes were due to adaptations in mitochondrial content or ETC protein abundance, as the abundance of citrate synthase (Figures 5E & 5F) was not different in any of the groups. Furthermore, PGC1-a mRNA abundance (Supplementary Figure 5A), and ETC complexes I, II, and IV, and ATP synthase protein abundance (Supplementary Figures 5B-E) were not significantly different between strains or diets.

**Figure 5.**
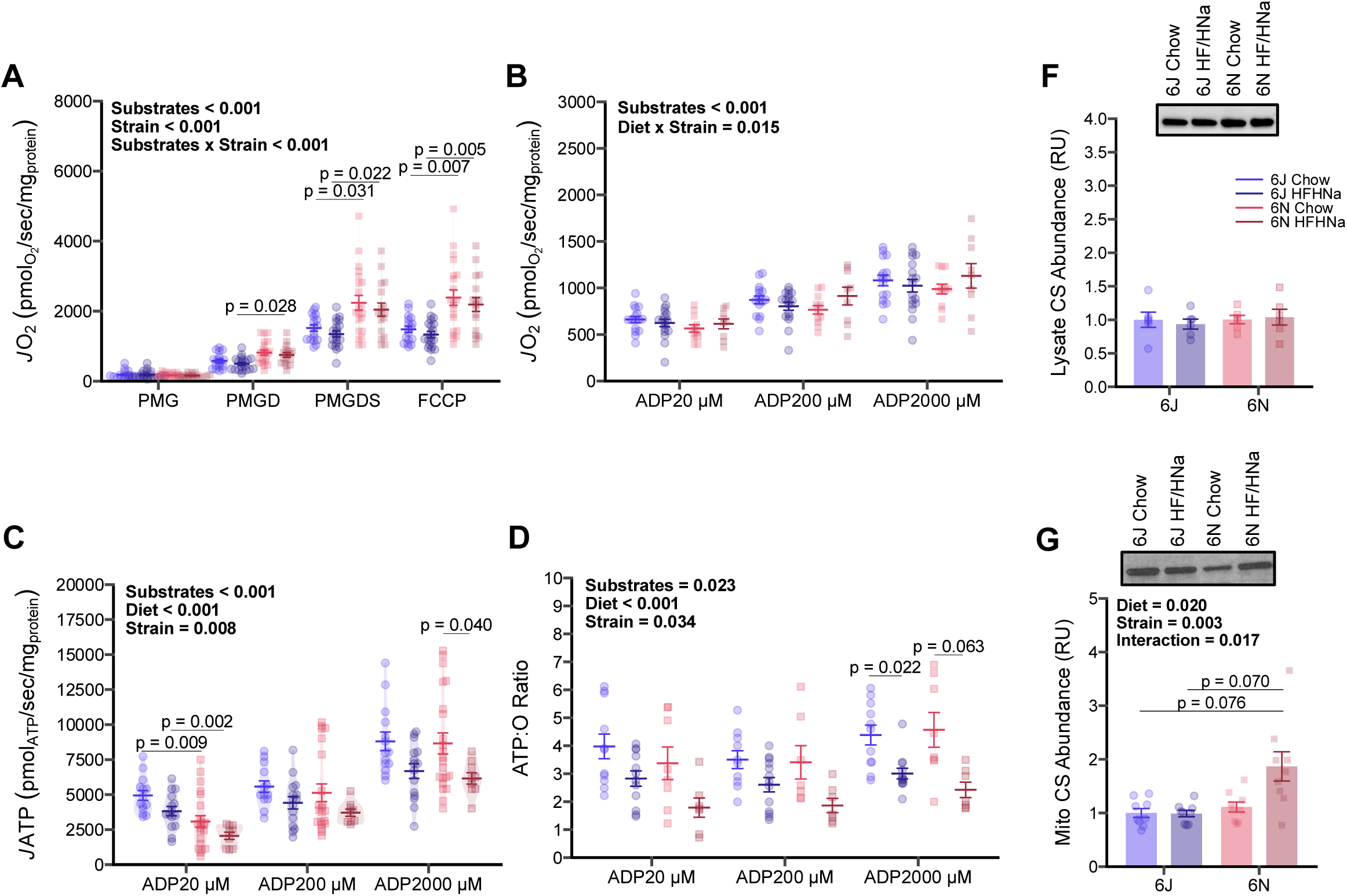
Mitochondrial respiration (*J*O_2_) in isolated mitochondria was not impacted by diet following a maximal (A) or submaximal (B) protocol. However, ATP production (*J*ATP; C) was significantly impaired in the mice fed a high-fat, high-salt (HFHNa) diet. Consequentially, mitochondrial ATP-to-O ratio (P:O Ratio; C) was decreased in the HFHNa-fed mice. These differences occurred despite no changes in mitochondrial citrate synthase (CS) protein abundance in whole-tissue lysate (F) or isolated mitochondria (G) preparations.

### Urine Proteomics

Last, preliminary analysis of urine proteins in 6J mice (chosen due to the diet-induced decline in GFR) revealed several urinary markers of interest that were increased with HF/HNa feeding (Figure 6A). Consistent with our urinalysis markers, albumin, cystatin C, and Lcn2 levels were increased in the urine of HF/HNa-fed mice, but the increase was not statistically significant, likely due to sample size limitations (Figure 6B). Interestingly, other kidney injury markers, such as Villin-1 (a structural protein found in tubular microvilli), were also increased in urine. Markers of proteostasis, such as the subunit c of the V0 domain of vacuolar H^+^-ATPase (Atp6v0c), sinal peptide peptidase-2A (Sppl2a), and programmed cell death 6 (Pdcd6), were increased in HF/HNa-fed mice urine. Likewise, several transport proteins, such as succinate dehydrogenase A (Sdha), aldolase B (Aldob), and the organic ion transporter (Slc22a19) were also increased in the urine. Collectively, these support several novel markers of interest to track the severity of renal failure in mice.

**Figure 6.**
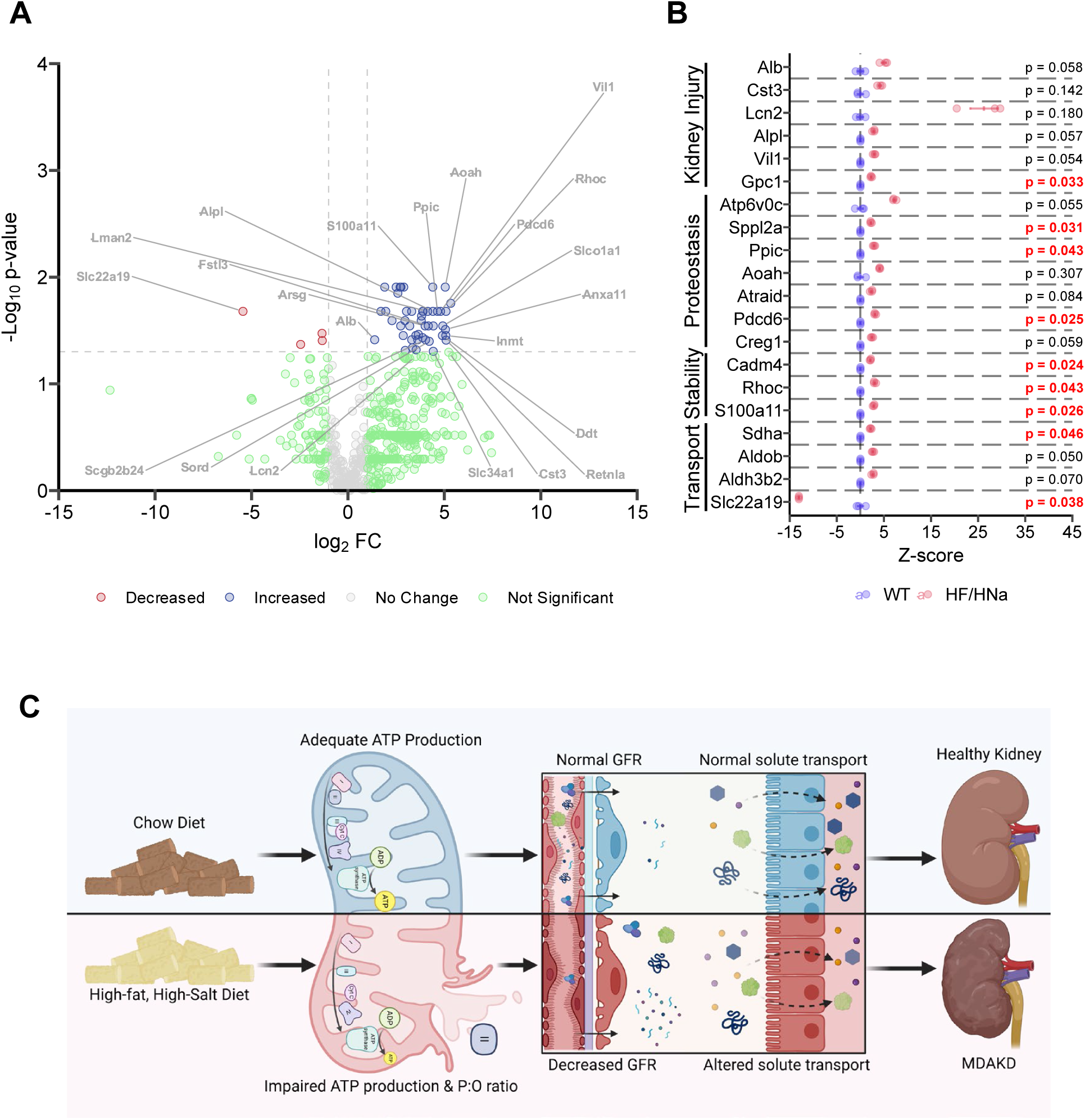
Urine proteomic analysis in a subset (n = 3 per group) of 6J mice (A) revealed that high-fat, high-salt (HFHNa) feeding significantly increased the presence of several urinary proteins. Several of these excreted proteins included those related to kidney injury, proteostasis, cell stability, and membrane transport (B). Collectively (C), our data support the hypothesis that a HFHNa diet induces impairments in mitochondrial ATP production, which drives decreases in glomerular filtration rate (GFR) and altered solute transport, leading to metabolic dysfunction-associated kidney disease (MDAKD).

## Discussion

In this study (summarised in Figure 6C), we demonstrate that 16 weeks of HF/HNa feeding in C57Bl/6 mice recapitulates several key features of MDAKDassociated CKD. Despite no effect on body weight or adiposity, HF/HNa-fed animals exhibited altered feeding behavior, increased water intake, and higher urinary output, consistent with prior reports that sodium-rich diets impose systemic metabolic stress independent of obesity.^12,18,19^ Functionally, 6J mice developed a mild but significant reduction in GFR, while 6N mice maintained or slightly increased GFR, reflecting strain-dependent susceptibility to renal injury. Both strains, however, displayed biochemical evidence of tubular stress with elevated U-cystatin C, U-KIM-1, and U-Lcn2, aligning with human CKD, where tubular injury markers often rise before overt albuminuria.^6,8,14,17,20^ At the molecular level, transporter expression patterns diverged between strains: 6N mice exhibited trends toward adaptive regulation of AQP2 and SGLT2, while 6J mice showed no changes in protein abundance or fibrotic gene expression, consistent with prior evidence that sodium transport and renal stress differ by genetic background.^30^ Finally, mitochondrial analyses revealed that HF/HNa feeding impaired ATP production and coupling efficiency without altering respiratory chain protein abundance, indicating functional bioenergetic uncoupling, in line with accumulating evidence that mitochondrial dysfunction is a central mediator of CKD progression.^16,21,23,25,28,41^ Further, we did not find any evidence of activation of pro-inflammatory or ER stress pathways, suggesting that mitochondrial bioenergetic deficiency is a significant contributor to early progression of renal dysfunction, regardless of genetic background. Together, these results establish HF/HNa feeding as a multifactorial model of CKD and highlight strain-specific adaptations that may underlie resistance versus susceptibility to diet-induced kidney dysfunction.

### Genetic Background Influences Renal Pathological Progression

A key novel finding of this study is the divergence between C57Bl/6J and C57Bl/6N mice in their renal response to HF/HNa feeding. While 6J mice exhibited a modest but significant decline in GFR and structural remodeling of glomeruli, 6N mice maintained or even slightly increased GFR despite equivalent dietary exposure. This contrast underscores the profound influence of genetic background on CKD susceptibility, consistent with prior work showing that allelic differences between C57 substrains shape cardiovascular, renal, and metabolic phenotypes.^30^ In all mice, lack of diet-induced glomerular morphological changes suggests that other factors, such as early hemodynamic impairment, rather than glomerulosclerosis, drive the initial loss of filtration capacity. Such a dissociation between functional decline and fibrotic remodeling closely mirrors early-stage human CKD, where reductions in GFR often precede substantial structural injury and where fibrosis becomes detectable only at later stages of progression.^1,3,13^

The 6N substrain, in contrast, exhibited relative preservation of filtration despite biochemical evidence of tubular stress, highlighting potential protective adaptations at the glomerular and vascular level. This observation is particularly important given that murine models often fail to develop measurable declines in GFR in response to dietary stress alone, limiting their translational relevance.^7,10,12–14,16,19,20,41^ Our results therefore establish the 6J strain as uniquely susceptible to hemodynamic perturbation under combined high-fat and high-salt feeding, while the 6N strain provides a valuable comparator in which protective adaptations can be studied. One possibility is that 6N mice exhibit more favorable autoregulatory control of renal blood flow, buffering glomeruli against hypertensive and osmotic stress. Another is that differences in sodium transporter regulation reduce the metabolic burden imposed on glomerular hemodynamics in 6N mice. By contrasting these two substrains, our study highlights the utility of genetic background as a tool to dissect mechanisms of susceptibility versus resilience, paralleling the heterogeneity observed in human CKD populations where some individuals remain resistant to renal decline despite comparable metabolic or hypertensive stressors.

### Tubular Stress is a Common Feature of HF/HNa Diet Feeding Regardless of Strain

Another important novel aspect of our study is the demonstration that tubular stress and transporter regulation diverge between strains, potentially explaining the preserved filtration capacity in 6N mice. Both substrains exhibited biochemical evidence of tubular injury, as shown by elevations in urinary cystatin C, KIM-1, and NGAL. These markers are increasingly recognized as early indicators of CKD progression in humans, often rising before overt albuminuria or reductions in eGFR are detectable.^6,13,44^ However, the molecular responses at the level of tubular transporters were strikingly strain-dependent. In 6N mice, AQP2 expression trended upward while SGLT2 trended downward, changes that would be expected to reduce osmotic stress by limiting glucose reabsorption while maintaining water handling. By contrast, 6J mice did not exhibit these adaptations.

These findings support a model in which protective adaptations in sodium and glucose transport in 6N mice reduce the energy burden of reabsorption and buffer glomeruli against hemodynamic overload. The absence of significant changes in ER stress markers (BiP, CHOP, PERK) and fibrotic and inflammatory transcripts (Col1a1, Col3a1, NF-κB, IL-6, CCL2, SGK1) in both strains suggests that these adaptations occur independently of canonical unfolded protein responses or pro-inflammatory pathways. Taken together, these results align with the hypothesis that tubular dysfunction and transport burden are central drivers of CKD progression and suggest that the balance between adaptive and maladaptive transporter regulation determines whether GFR is preserved or declines under chronic dietary stress.

### Mitochondrial Energetics as a Driver of Renal Stress

A central novel finding of this study is that HF/HNa feeding impaired renal mitochondrial energetics in both C57Bl/6 substrains, providing a unifying mechanism linking tubular transport burden to kidney injury. High-resolution respirometry demonstrated that while oxygen consumption rates (*J*O₂) were not significantly altered by diet, ATP production (*J*ATP) and the ATP:O ratio were reduced, consistent with bioenergetic uncoupling. This functional impairment occurred without significant changes in mitochondrial content, as assessed by citrate synthase abundance, or in respiratory chain complex protein levels, indicating that the observed deficits reflect bioenergetic dysfunction rather than reduced mitochondrial mass. Such uncoupling has been described as an early event in CKD pathogenesis, where increased sodium transport demand drives ATP turnover beyond the capacity of mitochondria to maintain efficient oxidative phosphorylation ^16,21,23–28,41,45^

The implications of these findings are twofold. First, they suggest that mitochondrial inefficiency may be a primary causative mechanism of HF/HNa-induced renal injury, independent of overt ER stress, fibrosis, or inflammation. Even modest reductions in ATP yield per oxygen consumed can compromise tubular reabsorptive function and contribute to the release of tubular injury markers, all of which were observed in this study. Second, the similar bioenergetic impairment in both strains suggests that mitochondrial uncoupling represents a shared pathogenic mechanism, while strain-specific differences in transporter regulation and glomerular hemodynamics determine whether GFR is preserved (6N) or declines (6J). These results align with recent reviews proposing mitochondrial dysfunction as a central mechanism of CKD progression^16,21,23,25^ and highlight mitochondrial energetics as a therapeutic target in diet-induced kidney disease.

### Conclusion & Future Directions

In conclusion, 16 weeks of HF/HNa feeding in C57Bl/6 mice reproduces several key features of MDAKD-associated CKD, including tubular injury, glomerular remodeling, and mitochondrial uncoupling, while revealing substrain-dependent susceptibility to filtration loss. These findings establish HF/HNa feeding as a tractable, non-surgical model of early diet-induced kidney disease and highlight the central role of mitochondrial energetics in driving renal stress. Future studies should expand on these findings by applying complementary approaches to dissect bioenergetic regulation at higher resolution,^46,47^ including measurements of mitochondrial membrane potential, reactive oxygen species production, and respiratory control in isolated nephron segments.^24,48^ In addition, comprehensive lipidomic profiling of mitochondrial phospholipids, particularly cardiolipin and phosphatidylethanolamine species, will provide critical insights into how changes in the mitochondrial membrane environment contribute to uncoupling and impaired ATP synthesis.^49–53^ Such studies will clarify the mechanistic links between tubular transport burden, mitochondrial dysfunction, and nephron loss, and may identify new therapeutic targets to preserve bioenergetic efficiency in CKD.

## Data Sharing

Data are available upon request. Scripts, training images, and output data are publicly available and accessible on GitHub at https://github.com/stdecker.

## Acknowledgments

We gratefully acknowledge the support of the University of Utah Health Sciences Core Facilities. Specifically, we thank the Cell Imaging Core Facility for support with microscopy, ARUP histological services, and the Genomics Core Facility for quantitative PCR services. Their expertise and resources were instrumental in the completion of this study. Schematics in this manuscript were generated using BioRender.

## Author Contributions

S.D., A.K., N.R., and K.F. conceived the project. S.D., Z.T.S., P.C.O., V.L.P., K.N.M, D.S., A.S.K., and S.R. performed experiments and collected data. S.D., Z.T.S., V.L.P., and D.S. analyzed the data. All authors contributed to the study design and data interpretation. S.D. and K.F. wrote the manuscript. All authors revised the manuscript and approved the final version.

## Supplementary material

Supplementary material is available online at www.kidney-international.org.

---

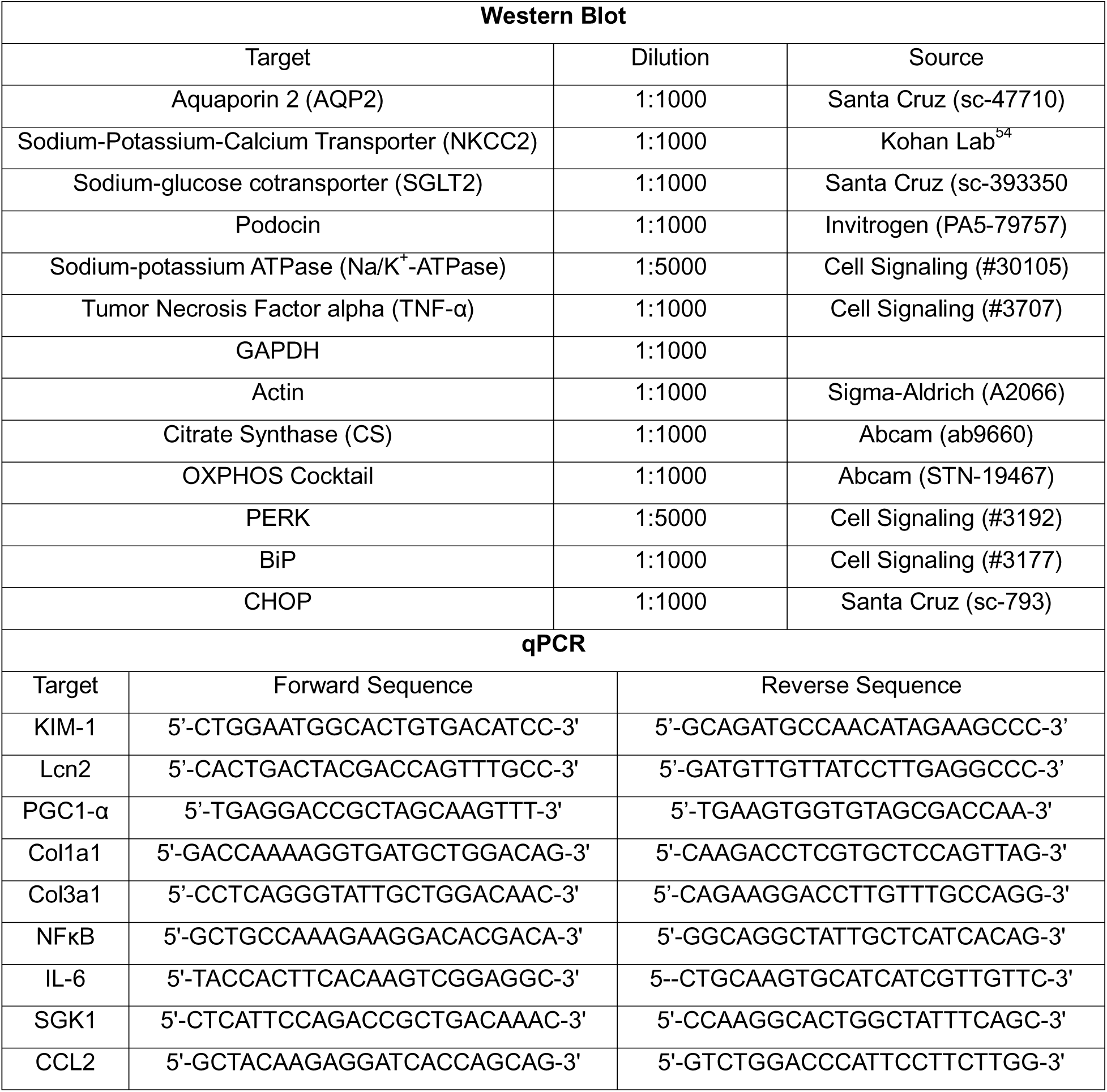

