## Supplemental Methods and Supplemental Figures S1-S5 for "A High-fat, High-salt Diet Model of MDAKD Impairs Bioenergetic Efficiency for ATP Synthesis"

15 N 2030 E

University of Utah

Salt Lake City, UT 84112

Keywords: Mitochondria, Metabolism, High-Salt, High-Fat, MDAKD

*Histology Analysis*

Trichrome-stained kidney sections were analyzed in Fiji software[33] (version 2.7.0) using a combination of custom macros, the Labkit plugin[34] (version 0.4.0), and a custom-made Python script to segment images using the MONAI framework[35-37] in a Google Colab environment. Images were converted in Fiji from .czi to .tif format using the BioFormats plugin. A subset of images was randomly selected and concatenated to be used as training images. A subset of glomeruli, tubules, hylum, and blood vessels was randomly selected and annotated in Labkit. Background and artifact features were also annotated to remove artifacts and improve learning. A deep-learning script in the MONAI framework was developed and implemented in Google Colab, which identified glomerular and tubular structures. This learning algorithm was used to segment glomerular and tubular structures and generate mask image files in the rest of the slides. Following segmentation, the generated masks were used to crop individual glomeruli and tubular structures for analysis. Analysis was performed using feature (i.e., size, circularity, etc.) cutoffs and the colour deconvolution2[38-40] (version 2.1) plugin incorporated into a custom macro in Fiji. Training images, masks, custom macros, and associated files are publicly available on Open Science Foundation (https://osf.io/user/fky4t) and GitHub (https://github.com/stdecker).

*Mass Spectrometry-based Proteomics*

Chemicals

LC-MS-grade solvents and mobile phase modifiers were obtained from Honeywell Burdick & Jackson, Morristown, NJ (acetonitrile, isopropanol, formic acid, urea), Fisher Scientific, Waltham, MA (Peptide desalting columns), Sigma Aldrich, St. Louis, MO (Tris-HCl, Dithiothreitol (DTT), Iodoacetamide (IAA)), Promega, Madison, WI (Sequencing grade trypsin/LysC), SPEX, Metuchen, NJ (Trifluoroacetic acid (TFA)).

Sample Preparation

Protein from ~200 µL of urine was precipitated with 9x volume of ice cold acetone and resuspended in 1x S-TRAP buffer (50 mM TEAB pH 8.5 and 5% SDS). 100 uL of additional 1x S-TRAP buffer was added to each sample and the total protein content was quantified using the Pierce™ BCA Protein Assay Kit. 10 µg of protein from each sample was added to a library sample. 10 µg of protein from each experimental sample was diluted into 25 µL of 1x S-TRAP buffer and the protein was reduced with 20 mM DTT at 37°C temperature for 30 minutes. Samples were alkylated with 40 mM IAA at room temperature for 45 minutes in the dark and acidified with 55% phosphoric acid to a final concentration of 4.5% phosphoric acid. 330 µL of binding buffer (10% 100 mM TEAB [final] pH 7.5 with phosphoric acid, 90% MeOH) was added to each sample and the samples were vortex and added directly to the S-TRAP column. The columns were centrifuged at 4000 x g for 1 min room temperature. Each column was washed 3x with 150 µL of binding buffer and the columns were transferred to a new clean tube. 10 µg of protein sample was digested with 0.75 µg of trypsin for 4 hours at 47°C. The library sample were digested with 2.5 µg of trypsin. The peptides were eluted from the column with 40 µL of 50 mM TEAB pH 8.5 followed by 40 µL of 0.2% formic acid and 40 µL of elution buffer (50 mM TEAB pH 8.5 + 0.2% formic acid + 50 % acetonitrile). The 150 µg library peptide sample was fractioned into 8 parts using Pierce™ High pH Reversed-Phase Peptide Fractionation Kit (Thermo Fisher Scientific) according to the manufacturer's instructions. All peptide samples were dried to completion and resuspended in 300 µL of 0.1% TFA. All peptide samples were desalted using Pierce™ Peptide Desalting Spin Columns (ThermoFisher Scientific) according to the manufacturer's instructions and resuspended in 45 µL of 0.1% formic acid. 3 µL from each experimental peptide sample was combined into a pooled sample to create the pooled QC sample to monitor instrument performance.

Mass Spectrometry Analysis

Reversed-phase nano-LC-MS/MS was performed on a nanoElute 2 (Bruker Daltonics) coupled to a Bruker timsTOF Pro2 mass spectrometer equipped with a nanoelectrospray source.

200 ng of each sample was injected directly onto the liquid chromatograph reverse-phase ReproSil C18 150 mm x 0.15 mm nanocolumn (Bruker Daltonics) heated to 50°C. The peptides were eluted with a gradient of reversed-phase buffers (Buffer A: 0.1% formic acid in 100% water; Buffer B: 0.1% formic acid in 100% acetonitrile) at a flow rate of 0.5 µL/min. The LC run lasted for 45 minutes with a starting concentration of 5% buffer B increasing to 28% buffer B over 40 minutes, up to 32% buffer B over 2 minutes and held at 95% B for 3 minutes. The separation column is equilibrated with 4 column volumes at 800 bar following each run. The timsTOF Pro2 was operated in PASEF data-dependent acquisition (DDA) MS/MS scan mode to generate the custom peptide library. The TIMS section was operated with a 120 ms ramp time at a rate of 7.93 Hz and an ion mobility scan range of 0.6-1.4 V·s/cm2. MS and MS/MS spectra were recorded from 100 to 1,700 m/z. A polygon filter was applied to select against singly charged ions. The quadrupole isolation width was set to 3 Da. The mass spectrometer was operated in PASEF data-independent acquisition (DIA) MS/MS scan mode to analyze each experimental sample. The TIMS section was operated with a 120 ms ramp time at a rate of 7.93 Hz and an ion mobility scan range of 0.6-1.4 V·s/cm2. dia-PASEF window parameters were set to mass width of 25 Da and 47 mass steps/ cycle. MS and MS/MS spectra were recorded from 147 to 1,322.6 m/z.

Proteomics Data Analysis

The custom peptide library was created using Fragpipe v22.0 software (downloaded 12-7-24) against the uniprot_ref_mouse database (12-7-2023 version with 17,191 proteins). Protein abundances based on peak intensity were calculated using DIA-NN software version 2.0. An allowance was made for 1 missed cleavage following trypsin digestion. No fixed modifications were considered. The variable modifications of methionine oxidation, N-terminal acetylation and cysteine carbamidomethylation (IAA) were considered with a mass tolerance of 15 ppm for precursor ions and a mass tolerance of 10 ppm for fragment ions. The results were filtered with a false discovery rate of 0.01 for both proteins and peptides. A minimum of 1 unique peptide should be reported for all proteins identified.

**Supplemental Figure 1.** Whole-cage food intake (A) and fat intake (B) of the groups. Body composition in a subset of mice (n = 5-7 per group) via NMR was used to quantify absolute and relative fat mass (C & D), lean mass (E & F), and fluid mass (G & H).

**Supplemental Figure 2.** Urine Cystatin C-to-creatinine ratio (A) and kidney injury marker 1 (KIM-1)-to-creatinie ratio (B) were increased only in the 6N mice fed a high-fat, high-salt (HFHNa) diet, but lipocalin 2 (LCN2)-to-creatinine ratio (C) was increased only in the HFHNa-fed 6J mice.

**Supplemental Figure 3.** A representative image of the structures identified by the Masson's trichrome deep-learning MONAI-based framework (A). High-fat, high-salt (HFHNa) diet did not impact the relative or absolute fibrotic areas in the tubules (B & C). Likewise, mRNA abundance of Col1α1 (D) and Col3α1 (E) were not influenced by diet. Representative images of H & E, PAS, and Jones H & E stains are shown in panels F-H, while representative images of the Jones Silver deep-learning MONAI-based framework are shown in panels I and J. Mice fed the HFHNa diet had lower corpuscle (K) and lower absolute basement membrane-stained (L areas, but relative staining of the basement membrane (M) was not influenced by diet.

**Supplemental Figure 4.** Representative images of western blots are shown in panel A. A high-fat, high-salt diet (HFHNa) did not have an effect on the protein abundance of podocin (B), $Na/K^+$-ATPase (C), or TNF-α (D).

**Supplemental Figure 5.** mRNA abundance of PGC-1α (A), nor protein abundance of mitochondrial complexes I, II, IV, or ATP synthase (B-E) were not influenced by diet. Representative blots are shown in panels F and G.

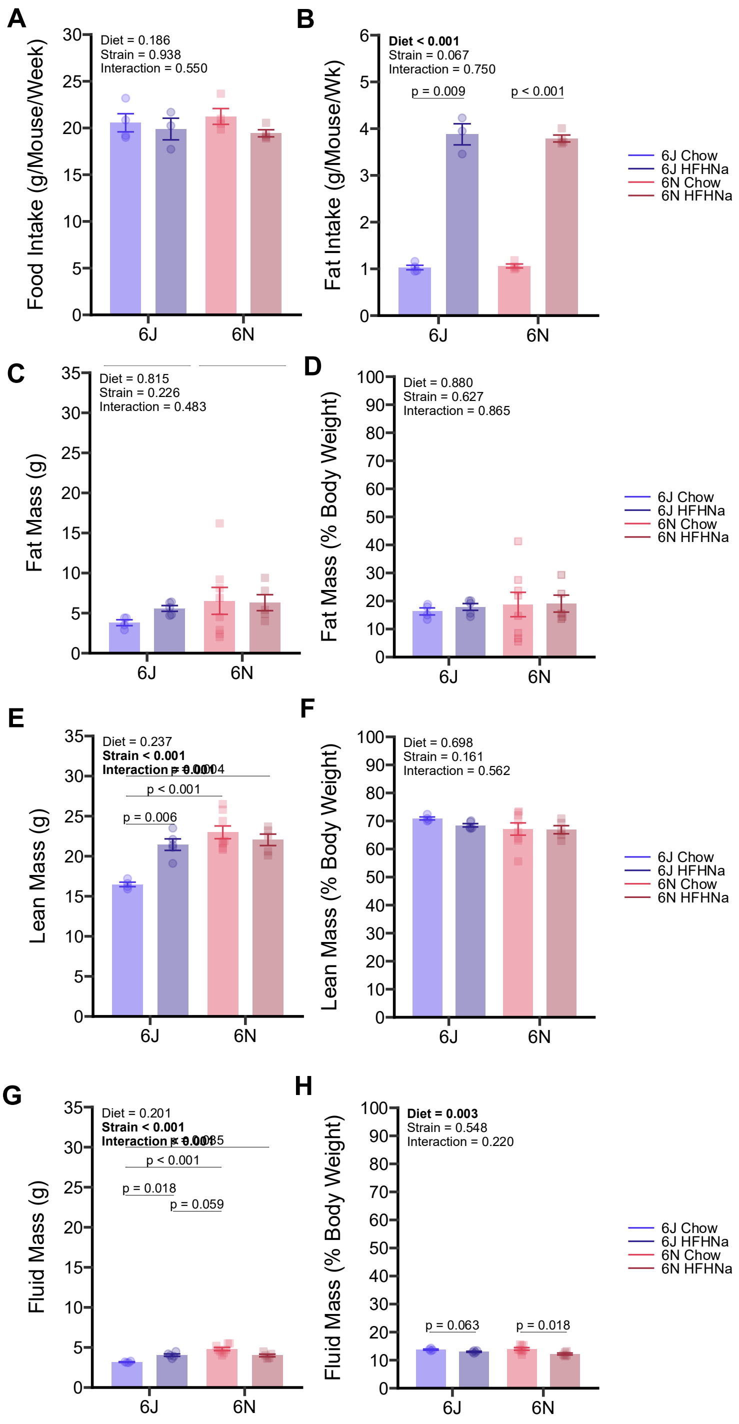
A
Food Intake (g/Mouse/Week)
Diet = 0.186
Strain = 0.938
Interaction = 0.550
6J
6N
B
Fat Intake (g/Mouse/Wk)
Diet < 0.001
Strain = 0.067
Interaction = 0.750
p = 0.009
p < 0.001
6J
6N
6J Chow
6J HFHNa
6N Chow
6N HFHNa
C
Fat Mass (g)
Diet = 0.815
Strain = 0.226
Interaction = 0.483
6J
6N
D
Fat Mass (% Body Weight)
Diet = 0.880
Strain = 0.627
Interaction = 0.865
6J
6N
6J Chow
6J HFHNa
6N Chow
6N HFHNa
E
Lean Mass (g)
Diet = 0.237
Strain < 0.001
p < 0.001
p = 0.006
6J
6N
F
Lean Mass (% Body Weight)
Diet = 0.698
Strain = 0.161
Interaction = 0.562
6J
6N
6J Chow
6J HFHNa
6N Chow
6N HFHNa
G
Fluid Mass (g)
Diet = 0.201
Strain < 0.001
p < 0.001
p = 0.018
p = 0.059
6J
6N
H
Fluid Mass (% Body Weight)
Diet = 0.003
Strain = 0.548
Interaction = 0.220
p = 0.063
p = 0.018
6J
6N
6J Chow
6J HFHNa
6N Chow
6N HFHNa

Figure S1

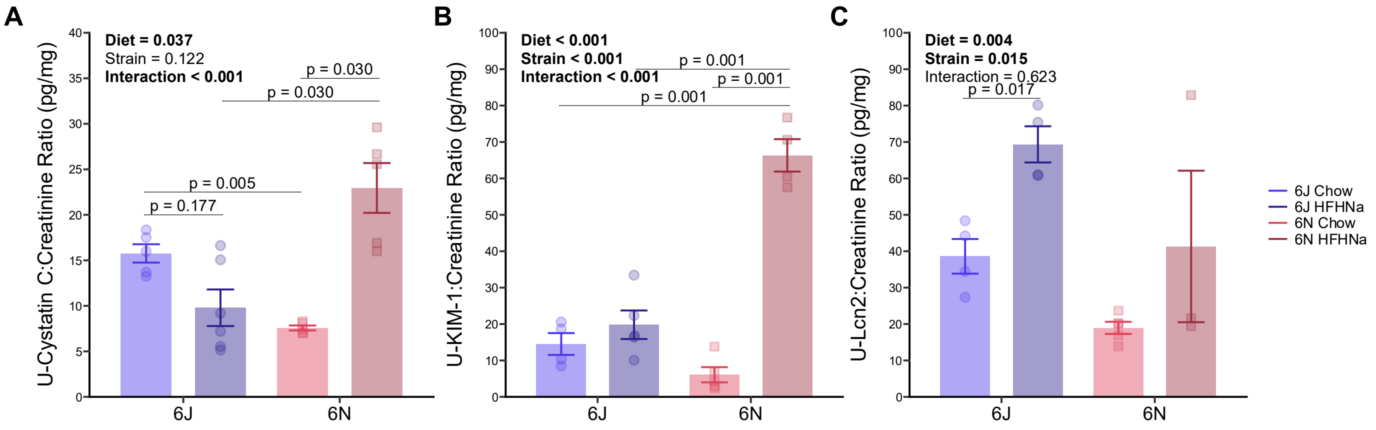
A
Diet = 0.037
Strain = 0.122
Interaction < 0.001
p = 0.030
p = 0.030
p = 0.005
p = 0.177
U-Cystatin C:Creatinine Ratio (pg/mg)
6J
6N
B
Diet < 0.001
Strain < 0.001
Interaction < 0.001
p = 0.001
p = 0.001
p = 0.001
U-KIM-1:Creatinine Ratio (pg/mg)
6J
6N
C
Diet = 0.004
Strain = 0.015
Interaction = 0.623
p = 0.017
U-Lcn2:Creatinine Ratio (pg/mg)
6J
6N
6J Chow
6J HFHNa
6N Chow
6N HFHNa

Figure S2

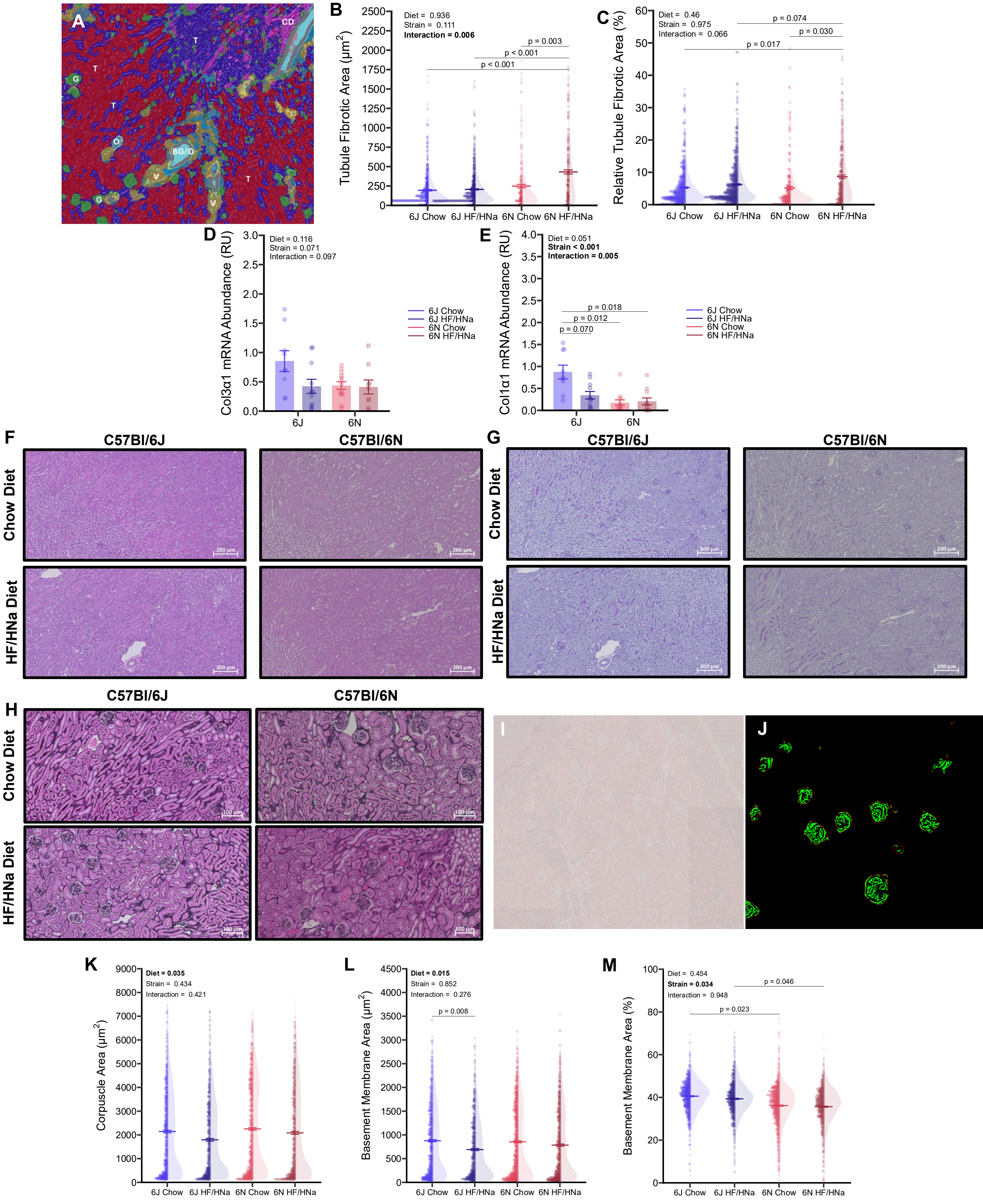
A
CD
T
G
T
T
O
BG/O
V
T
G
V
B
Diet = 0.936
Strain = 0.111
Interaction = 0.006
p = 0.003
p < 0.001
p < 0.001
Tubule Fibrotic Area (μm$^2$)
6J Chow
6J HF/HNa
6N Chow
6N HF/HNa
C
Diet = 0.46
Strain = 0.975
Interaction = 0.066
p = 0.074
p = 0.030
p = 0.017
Relative Tubule Fibrotic Area (%)
6J Chow
6J HF/HNa
6N Chow
6N HF/HNa
D
Diet = 0.116
Strain = 0.071
Interaction = 0.097
Col3α1 mRNA Abundance (RU)
6J
6N
6J Chow
6J HF/HNa
6N Chow
6N HF/HNa
E
Diet = 0.051
Strain < 0.001
Interaction = 0.005
p = 0.018
p = 0.012
p = 0.070
Col1α1 mRNA Abundance (RU)
6J
6N
6J Chow
6J HF/HNa
6N Chow
6N HF/HNa
F
C57Bl/6J
C57Bl/6N
Chow Diet
HF/HNa Diet
200 μm
G
C57Bl/6J
C57Bl/6N
Chow Diet
HF/HNa Diet
200 μm
H
C57Bl/6J
C57Bl/6N
Chow Diet
HF/HNa Diet
100 μm
I
J
K
Diet = 0.035
Strain = 0.434
Interaction = 0.421
Corpuscle Area (μm$^2$)
6J Chow
6J HF/HNa
6N Chow
6N HF/HNa
L
Diet = 0.015
Strain = 0.852
Interaction = 0.276
p = 0.008
Basement Membrane Area (μm$^2$)
6J Chow
6J HF/HNa
6N Chow
6N HF/HNa
M
Diet = 0.454
Strain = 0.034
Interaction = 0.948
p = 0.046
p = 0.023
Basement Membrane Area (%)
6J Chow
6J HF/HNa
6N Chow
6N HF/HNa

Figure S3

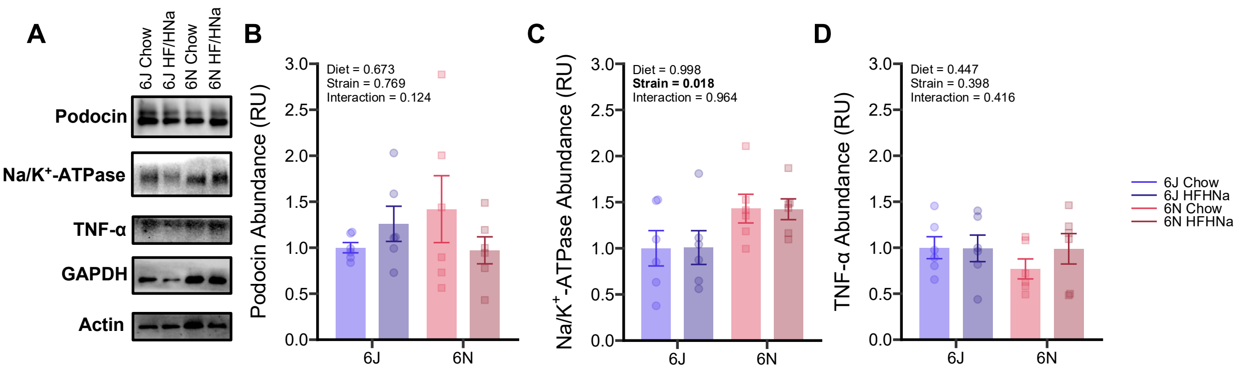
A
6J Chow
6J HF/HNa
6N Chow
6N HF/HNa
Podocin
Na/K+-ATPase
TNF-α
GAPDH
Actin
B
Diet = 0.673
Strain = 0.769
Interaction = 0.124
Podocin Abundance (RU)
C
Diet = 0.998
Strain = 0.018
Interaction = 0.964
Na/K+-ATPase Abundance (RU)
D
Diet = 0.447
Strain = 0.398
Interaction = 0.416
TNF-α Abundance (RU)
6J
6N
6J Chow
6J HFHNa
6N Chow
6N HFHNa

Figure S4

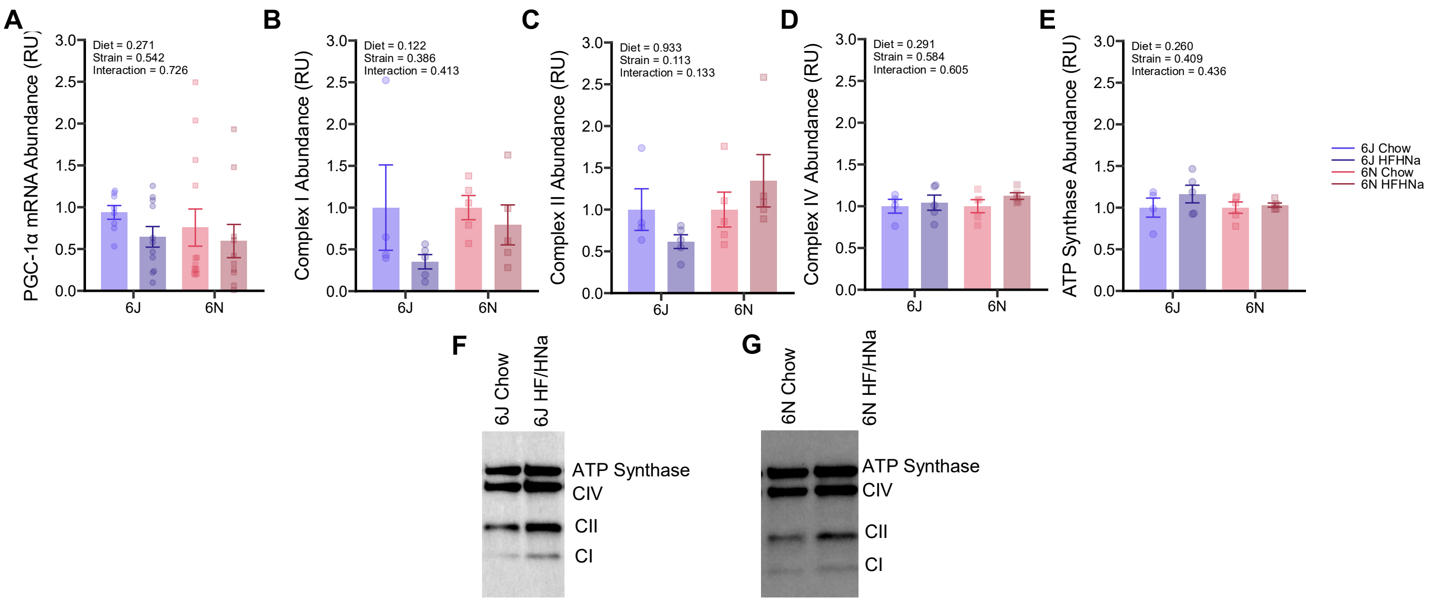
A
PGC-1α mRNA Abundance (RU)
Diet = 0.271
Strain = 0.542
Interaction = 0.726
B
Complex I Abundance (RU)
Diet = 0.122
Strain = 0.386
Interaction = 0.413
C
Complex II Abundance (RU)
Diet = 0.933
Strain = 0.113
Interaction = 0.133
D
Complex IV Abundance (RU)
Diet = 0.291
Strain = 0.584
Interaction = 0.605
E
ATP Synthase Abundance (RU)
Diet = 0.260
Strain = 0.409
Interaction = 0.436
6J
6N
6J Chow
6J HFHNa
6N Chow
6N HFHNa
F
6J Chow
6J HF/HNa
ATP Synthase
CIV
CII
CI
G
6N Chow
6N HF/HNa
ATP Synthase
CIV
CII
CI

Figure S5
